# Programmatic design and editing of *cis*-regulatory elements

**DOI:** 10.1101/2025.04.22.650035

**Authors:** Jacob Schreiber, Franziska Katharina Lorbeer, Monika Heinzl, Franziska Reiter, Baptiste Rafanel, Yang Young Lu, Alexander Stark, William Stafford Noble

## Abstract

The development of modern genome editing and DNA synthesis has enabled researchers to edit DNA sequences with high precision but has left unsolved the problem of designing these edits. We introduce Ledidi, a computational method that rephrases the discrete design task of choosing which edits to make as an easily solvable continuous optimization problem. Ledidi can use any pre-trained deep learning model to guide the optimization, yielding an edited sequence that exhibits the desired outcome while explicitly minimizing the number of edits used. When applied in dozens of settings, we find that Ledidi’s designs can precisely control transcription factor binding, chromatin accessibility, transcription, and enhancer activity *in silico*. By using several deep learning models simultaneously, we design cell type-specific enhancers and experimentally validate them *in cellulo*. Finally, we introduce the concept of an “affinity catalog”, where the design task is repeated multiple times across continuous variants of the design target. We demonstrate how these catalogs can be used to interpret deep learning models and the impact of starting template sequences, and also to design regulatory elements that control transcriptional dosage while maintaining cell type-specificity.

## 1 Main

Recent advances in genome editing, driven by the discovery and development of the CRISPR-Cas9 system [1–4] and subsequently base [5–7] and prime editing [8–10], have significantly reduced the cost and effort required to precisely modify genomic sequence within living cells. Moreover, improvements in DNA synthesis have enabled the creation of essentially any DNA sequence at high throughput and low cost, sparking unprecedented opportunities for synthetic biology. Consequently, genome editing is now routinely used in a variety of contexts, including engineering crops [11, 12], development of therapeutics [13, 14], and scientific investigation [15–19]. To avoid unintended consequences and potential off-target effects, a desirable strategy in these applications is to stay close to naturally occurring sequences by only making a relatively small number of edits to achieve the specific target function. However, despite these technical capabilities, choosing which edits to make to achieve these functions remains elusive.

In parallel, a surge in the amount and quality of genomics data [20–22] has led to the development of increasingly sophisticated genomic machine learning (ML) models [23, 24]. These models can make highly accurate predictions for transcription factor (TF) binding [25, 26], histone modification, chromatin accessibility [27–30], various measures of transcription [31–33], alternative splicing [34, 35], enhancer activity [36], and more. Although these models differ substantially in architecture and precise output, a commonality is that they are sequence-based, in that they learn regulatory patterns encoded in DNA sequence to make predictions. Because these models are now highly accurate across a variety of genomic modalities, there is an urgent need for approaches that make them available for design settings to enable synthetic biology.

Previous work has successfully used such ML models to design regulatory sequences *de novo*. In this paradigm, DNA sequences are proposed by a design algorithm (sometimes an ML model itself) and are evaluated by a predictive ML model (the “oracle”), with iterative updates occurring until the desired goal is achieved. This strategy has been used to design enhancers active in *Drosophila melanogaster* and human cell lines [36–38]. We recommend De Winter, Konstantakos, and Aerts [39] as a comprehensive review of DNA sequence design methods.

Although *de novo* design provides compositional flexibility, it is not always the most practical approach. For example, base and prime editors can make precise edits but perform best when only a few changes are required. Model-based design methods can only be as good as the underlying oracles, yet many oracles have difficulty generalizing too far from the reference sequences on which they were trained [40–42]. Consequently, designing DNA by starting with an initial template that exhibits many desirable characteristics, such as CpG or motif content, and then minimally altering it may be more robust than *de novo* design. Such a method would simplify quality control by focusing attention on the small number of nucleotides that are changing and their immediate context, e.g., presence within a TF binding site, and also enable scientific exploration by providing a lens into the learned *cis-*regulatory logic of the oracle model.

Accordingly, we propose Ledidi, a computational method for designing *edits* to an initial template sequence. Ledidi rephrases the discrete task of choosing which edits to make as a continuous optimization problem using the straight-through estimator [43]. In this formulation, the loss function being optimized is comprised of two components: the deviation from the target outcome and a novel input loss, which counts the number of edits being proposed. By explicitly penalizing the number of edits, Ledidi regularizes the optimization problem and implicitly limits off-target effects and unintended functional changes whose consequences may not be well captured by the oracle model. Despite this focus on edits, Ledidi can still make *de novo* designs, if necessary, by beginning its optimization from a randomly generated template. Across dozens of oracle models and diverse genomic modalities, we find that Ledidi can quickly generate compact sets of edits that are predicted to yield target outcomes.

As a validation, we design cell type-specific enhancers in *Drosophila melanogaster* cell lines and experimentally characterize them using Self-Transcribing Active Regulatory Region sequencing (STARR-seq, [44]), a massively parallel reporter assay (MPRA) that measures their transcriptional influence. We found that Ledidi’s designs exhibit strong cell type-specific signals and that many of the designs exhibit stronger regulatory activity than the strongest endogenous elements. These designs require a small number of edits and changed motif composition in anticipated ways. By inspecting these designs using feature attribution methods, we pinpoint the precise consequence of each edit and find that many edits played multiple roles, e.g., destroying a binding site in one cell type while also playing a role in creating one in the other, or being involved in overlapping binding sites in both cell lines.

Finally, we introduce the concept of an “affinity catalog”, where the design process is repeated across multiple levels of activity instead of only aiming for the strongest activity. When used as a tool for scientific investigation, affinity catalogs can be a powerful approach for distilling the learned *cis-*regulatory logic of the oracle models while also providing insights into latent binding sites in the starting template sequences, which frequently contain regions that are only a small number of nucleotide edits away from TF binding sites. When validated using an MPRA, we found that affinity catalogs designed by Ledidi control not only cell type-specificity but also transcriptional dosage in the on-target cell line.

## 2 Ledidi achieves functional outcomes with few edits

Ledidi designs edits to an initial template sequence, with the goal of achieving specified characteristics as predicted by one or more oracle ML models (Fig 1A). By focusing on the design of a small number of edits, we contrast with the more common *de novo* design setting, where entire sequences are created from scratch. Briefly, Ledidi works by learning an underlying weight matrix that is the same size as the template sequence, where the combination of these weights and the template is interpreted as a probability distribution over the nucleotides (see Methods for details). Sequences containing edits are sampled from these probability distributions and passed through the oracle model. The model’s predictions are compared to the desired output to calculate a loss, and the gradient of this loss is used to update the underlying weight matrix. In this formulation, Ledidi’s total loss is made up of two terms: the output loss, as described above, and a novel input loss term that counts the number of edits to the original sequence. This second loss term enables Ledidi to explicitly minimize the number of edits needed to achieve the desired objective.

**Figure 1:**
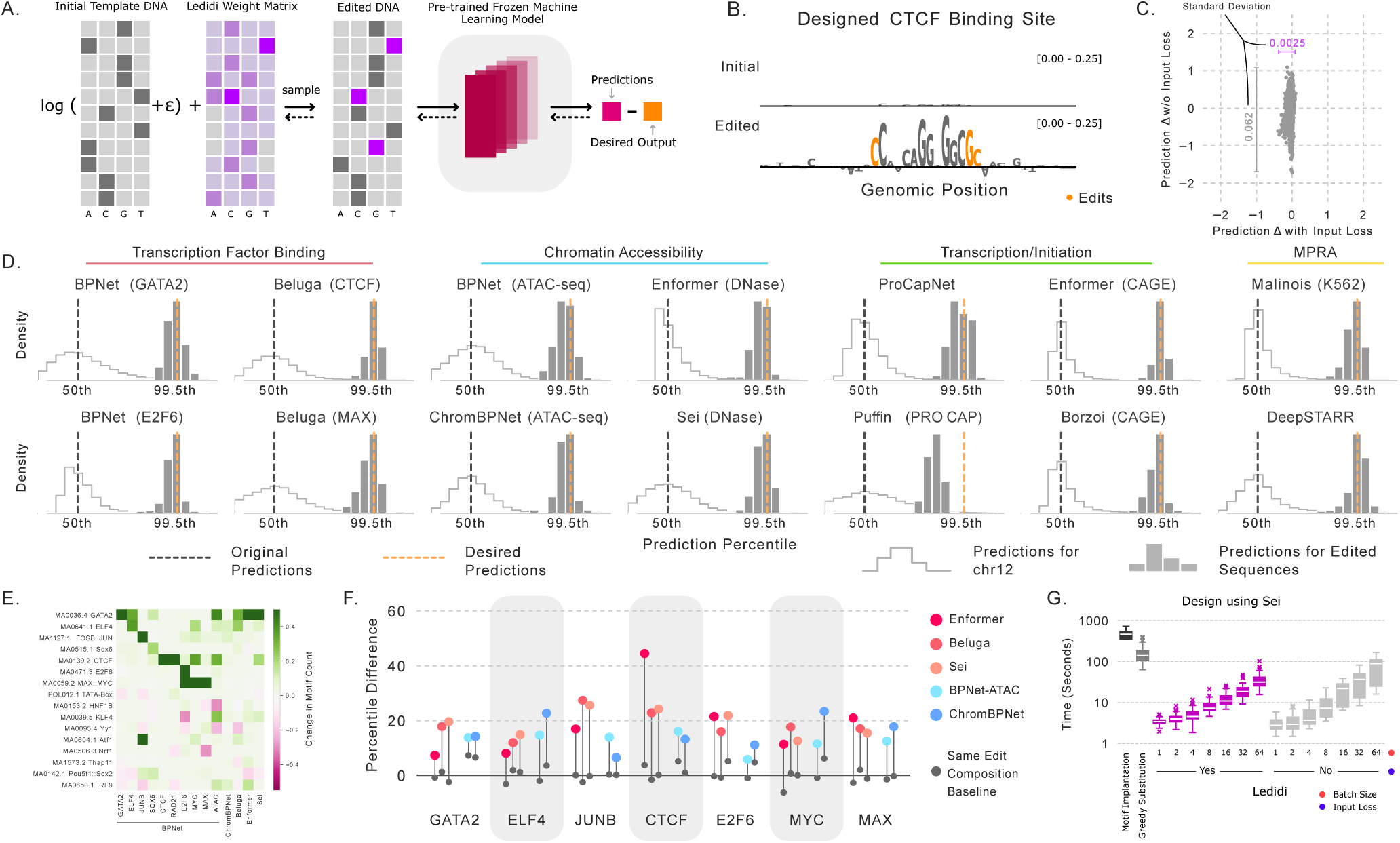
Ledidi accurately designs minimal DNA edits *in silico*. (A) A schematic of the Ledidi method, where edits are sampled by combining the template sequence and a learned weight matrix and evaluated using a pre-trained ML model. (B) An example of a designed CTCF site, where only a few edits (in orange) are necessary to create predicted binding. (C) Differences in prediction from Enformer, with each dot representing one task, when using or not using an input loss, with the standard deviation on each axis noted. (D) Examples of predictions for Ledidi-based designs in 14 settings covering four modalities. (E) The change in motif composition (rows) when designing strong sites for each model (columns). (F) External validation of Ledidi’s edits using predictions from models not involved in design. (G) Timings for various design methods across nine design tasks involving Sei.

As an initial evaluation, we pair Ledidi with a DeepSEA/Beluga oracle to design CTCF binding sites in templates where CTCF does not bind (see Methods for details). When tasked with significantly increasing binding, we find that Ledidi makes an average of 4.34 ±1.72 edits across 160 2 kbp designs (Supplementary Fig 1) with over 36% of cases needing only 3 edits. This number is significantly smaller than one might expect given the length of the CTCF motif (∼19 bp). When we more closely examined a representative example, we found that Ledidi was targeting latent binding sites that were almost a CTCF motif but had mismatches at critical positions in the corresponding position weight matrix (Fig 1B). By editing these critical nucleotides, Ledidi completed a CTCF motif using far fewer edits than would result from implanting an entire motif into the template. As we will see, this succinctness is not uncommon across design tasks and we frequently observed latent binding sites that, with only a small number of edits, became interpreted as TF binding sites by the oracles.

An important motivation for minimizing the number of edits is that this approach may yield designs with fewer unintended functional consequences. Accordingly, we next investigated the magnitude of these consequences by pairing Ledidi with BPNet to design GATA2 binding sites in a 2,114 bp long sequence with and without the penalty on making edits. As expected, penalizing edits resulted in designs with significantly fewer edits (10.9±4.4) than not penalizing them (687.9 ± 323.0). In one case, a sequence designed without an edit penalty contained 955 edits — redesigning almost half of the sequence. Making so many edits is not unexpected because, without an input loss, Ledidi has no reason to revert neutral edits that arise through the stochastic sampling process. However, these edits substantially changed the GC content (40.3% to 46.0%), whereas penalizing edits kept GC content mostly the same (40.3% to 40.5%). To better understand the global implications of these edits, we used a different ML model, Enformer, to compare the functional profiles of these loci in human cells before and after making edits. Considering all 5313 genomic functions predicted by Enformer, we found that penalizing edits resulted in much more similar predicted profiles globally (Fig 1C, *r* = 0.9912 with penalty, *r* = 0.8566 without). This finding demonstrates that designing sequences solely with a focus on one characteristic may inadvertently lead to substantial changes in other aspects, even if the design target is something conceptually simple like inducing GATA binding. However, Ledidi’s strategy of penalizing edits can mitigate these effects.

## 3 Ledidi is a general-purpose designer of edits

Unlike generative models for design, Ledidi does not require the training and validation of a new ML model for each design setting, but rather can be paired with any sequence-based ML model. To demonstrate this flexibility, we evaluated how well Ledidi can design edits out-of-the-box across a wide range of genomic modalities and predictive models, most of which were not trained by us. Specifically, we considered 14 design settings, including TF binding, chromatin accessibility, transcription, and MPRA activity, using ten different oracle models (BPNet [26], Beluga [45], ChromBPNet [27], Enformer [32], Sei [29], ProCapNet [31], Puffin [46], Borzoi [33], Malinois [38] and DeepSTARR [36], Supplementary Table 1). To account for technical differences in the experimental readouts and other differences in post-processing when training these models, we phrased each design problem as editing an initial template exhibiting median activity into one exhibiting 99.5th percentile activity (see Methods for details).

We observed that Ledidi’s designed edits dramatically increased predicted activity in each of the 14 settings (Fig 1D). On average, Ledidi made 16.54 edits per sequence, driving the predicted activity from the 50th to the 99.27th percentile. Puffin was an outlier, requiring 69.09 edits on average yet only yielding predictions in the 97.54th percentile, whereas Enformer (CAGE) and DeepSTARR yieldied the best designs, at the 99.48th percentiles, using averages of 13.83 edits in Enformer’s 2 kbp input window and 11.08 edits in DeepSTARR’s 249 bp input window, respectively. Beluga required the fewest edits, with only 4.34 edits in a 2 kbp input window to bring CTCF binding to the 99.45th percentile and 9.07 edits to bring MAX to the 99.39th percentile. We found that the edits created motif instances known to be associated with the predicted genomic modalities, e.g., the motif associated with the TFs whose binding is being predicted, with several interesting co-binding relationships emerging from the designs (Fig 1E). Together, these findings show that Ledidi is a general purpose designer of edits, but the number of edits proposed is influenced by the accuracy of the model and the complexity of the design goal.

A potential concern when performing model-based design is that the designs may be overly reliant on learned statistical shortcuts [47] or spurious correlations [48] encoded in the oracle model. Accordingly, we evaluated whether binding sites designed using BPNet models for seven TFs elicited responses in matched outputs from models not involved in the design process (Fig 1F, see Methods for details). These external models included Enformer, Beluga, and Sei, which predict matched TF binding directly, and BPNet-ATAC and ChromBPNet, which predict chromatin accessibility that we expect will increase as a consequence of TF binding. We found that Ledidi’s edits increase the predictions from these models by an average of 16.9 percentiles, with CTCF edits yielding the strongest response (+24.2 percentiles), and JunB edits yielding the weakest (+14.5 percentiles). These averages are driven by a strong response to CTCF edits by Enformer (+44.4 percentiles) and a particularly weak response from ChromBPNet to JunB edits (+6.5 percentiles). In contrast, when we apply the same *composition* of edits (i.e., number of A → C and T → G mutations, etc.) randomly to the initial sequences, we find only a 0.7 percentile increase on average. In each setting, this baseline was significantly weaker than the designed edits, further demonstrating that Ledidi’s edits are not based on shortcuts and do more than simply change nucleotide composition, e.g., GC content.

Finally, an advantage of gradient-based design methods like Ledidi is that they can be significantly faster than prediction-based design methods. As a demonstration, we compared the speed of Ledidi against two common prediction-based design methods: motif implantation and greedy substitution (sometimes called “directed evolution”). Across nine design tasks that use Sei — a multi-task model that makes predictions for 21,907 forms of activity — we find that Ledidi with its default batch size of 16 was 40.7x faster than motif implantation and 14.1x faster than greedy substitution (Fig 1G). These speedups match our intuition because gradient-based methods replace repeated forward-based passes per round with a single backward pass. Although the input loss constrains the design task and could hypothetically lead to faster convergence, we only noticed a small speedup with an input loss and only when using larger batch sizes.

## 4 Designing cell type-specific enhancers

Having demonstrated that Ledidi can design edits driving a diverse range of transcriptional and epigenomic responses, we next tackled the more challenging task of designing elements that exhibit cell type-*specific* activity. Here we build upon previous work from de Almeida *et al.* [36, 37] and Taskiran *et al.* [49], who designed enhancers active in a target cell type or tissue, to design enhancers whose activity is now specific to a target cell type. This task is more challenging because it requires that the designs must avoid *cis-*regulatory features that drive activity in both cell types.

Our designs targeted the Schneider-2 (S2) and ovarian somatic cell (OSC) types of *Drosophila melanogaster*, and we used five pools of initial templates: genomic sequences inactive in both cell types, active only in S2 cells, active only in OSC cells, active in both cell types, and uniformly randomly generated sequences (see Methods for details, Fig 2A). We paired Ledidi with two DeepSTARR models that had been independently trained in each cell type, and we optimized a modified objective that used the min-gap loss proposed by Gosai *et al.* [38] for the output loss (Fig 2B). This loss attempts to maximize the margin in predictions between the on- and off-target cell types, without trying to hit specific target values. For each pool, we tasked Ledidi with editing the template to be S2- or OSC-specific. We found that the predictions for the designed elements were strongly cell type-specific, with a slight preference for regions that began as enhancers in the target cell type versus regions that began as enhancers in the other cell type (Fig 2C).

**Figure 2:**
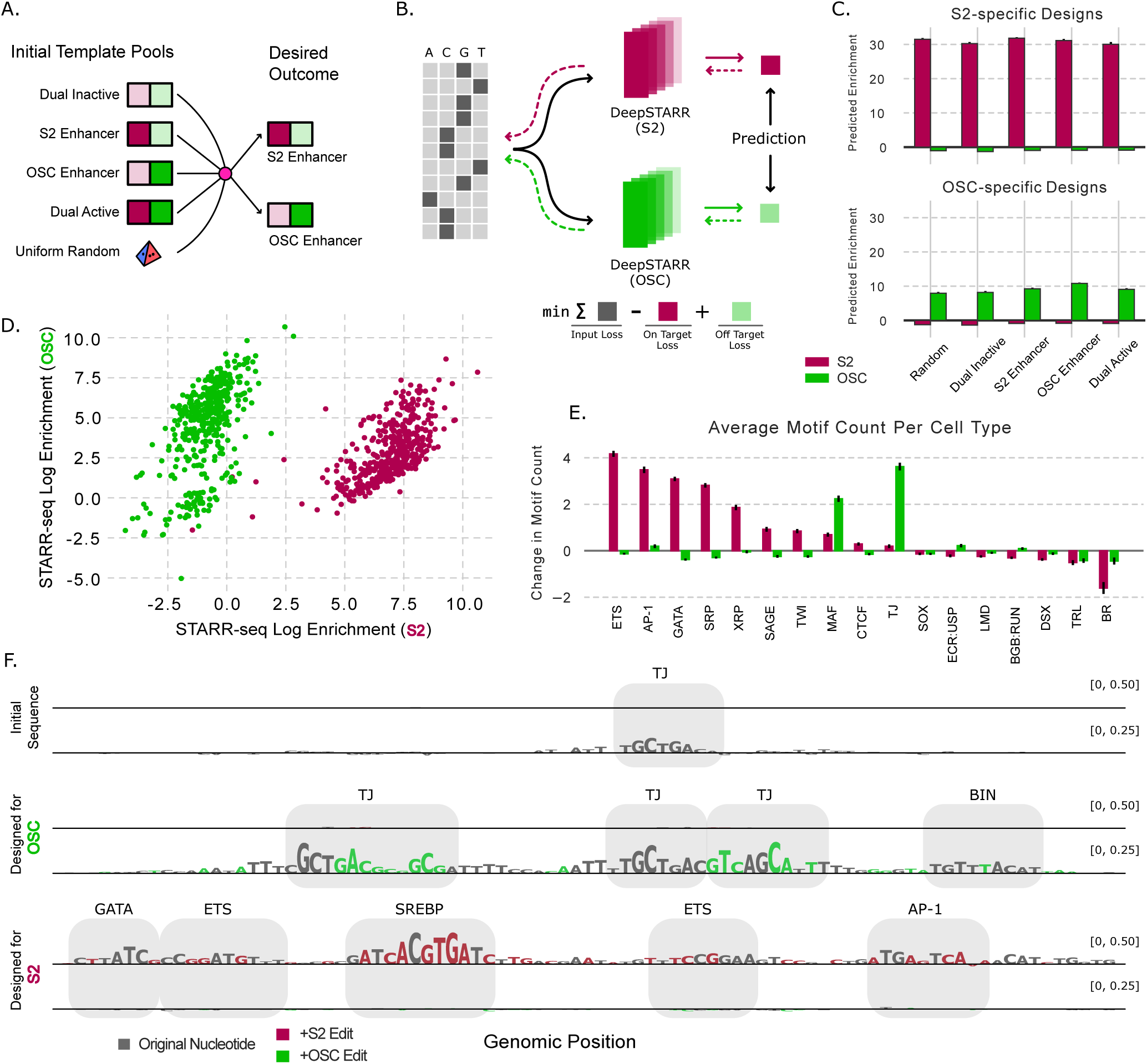
Design and validation of cell type-specific enhancers. (A) A cartoon visualizing the design process as taking genomic sequences from several pools and editing each into either an S2- or an OSC-specific enhancer. (B) A schematic demonstrating how multiple DeepSTARR models are used simultaneously by Ledidi to design edits using a min gap-based loss. (C) Predictions from the DeepSTARR models on the designed S2-specific enhancers (above) and OSC-specific enhancers (below). (D) Experimental STARR-seq data on the enhancers designed to be S2-specific (in burgundy) or OSC-specific (green). The lower left region of the plot corresponds to enhancers that were not expressed in either cell type. (E) The change in motif count after incorporating the proposed edits. (F) Attributions from both DeepSTARR models on an exemplar sequence that was originally an OSC-specific enhancer, and edited versions making it a stronger OSC enhancer or an S2-specific enhancer.

We then experimentally validated the designs using a STARR-seq experiment (see Methods for details) and observed that the designs generally exhibited strong and specific activity (Fig 2D). Elements designed to be OSC-specific (in green) exhibited a median enrichment in transcript abundance over the input control of 38.65x in OSC cells but only a 0.59x enrichment in S2 cells (65.5x stronger enrichment in the on-target cell line), and elements designed to be S2-specific (in burgundy) exhibited a 135.8x enrichment in S2 cells and a 6.36x enrichment in OSC cells (21.4x stronger enrichment in the on-target cell line). All initial template pools could be edited to become cell type-specific enhancers of each cell type, even when this meant swapping the cell type the element is active in (Supplementary Fig 2). Further, 419 out of 448 (93.5%) of the designed S2-specific enhancers and 72 out of 423 (17.0%) of the designed OSC-specific enhancers exhibited stronger activity than the average endogenous enhancer in the strongest bin of a ladder designed to benchmark activity (see Methods for details). 42 (10.7%) of the designed S2-specific enhancers and one of the designed OSC-specific enhancers exhibiting activity stronger than the strongest endogenous enhancer, demonstrating that Ledidi can design enhancers outside the scope of the training data used for the underlying DeepSTARR models.

However, two issues arose from the use of the min-gap loss. First, because this loss only maximizes the margin between the two cell lines without requiring that the off-target cell line exhibits near-zero activity, many of the S2-targeted designs exhibit activity in both cell lines, albeit much stronger in S2 cells. Second, a population of OSC-targeted designs exhibit no activity in either cell line. In this population, the predictions from the OSC DeepSTARR model are significantly weaker than the other OSC-specific enhancers but still stronger than the S2 predictions (Supplementary Fig 3). These results suggest that future efforts should aim to supplement the min-gap loss with an additional loss term requiring the off-target cell line to exhibit predicted activity of endogenous non-enhancer regions.

Next, we considered the changes to motif composition induced by Ledidi’s edits when designing these regulatory elements. Across all template pools, we found that Ledidi made 100.4±9.4 edits when designing S2-specific elements and 53.8±16.1 edits when designing OSC-specific elements given a 249 bp template (Supplementary Fig 4). These edits significantly increased the number of motifs commonly associated with ETS, AP-1, and GATA binding for the S2-specific elements but left the count of these motifs mostly unchanged in the OSC-specific elements (Fig 2E). In contrast, traffic jam (TJ) and MAF motifs showed the opposite patterns. More broadly, the enriched motifs for both cell types are consistent with those known to be important [50] (Supplementary Fig 5). Curiously, BR showed a decrease in motif count in both cell lines, and further investigation suggests that this may not be because BR is a repressor but, rather, that it is similar to AP-1 motifs and can be edited into those.

Finally, we more closely inspected the edits proposed by Ledidi at a representative locus that was initially an experimentally verified OSC-specific enhancer (Fig 2F). DeepSTARR model attributions reveal that a TJ motif is the primary driver of predicted activity. When edited to become a stronger OSC enhancer, Ledidi makes edits that add additional TJ motifs to the surrounding area. However, when the same region is edited to become an S2-specific enhancer, two things happen. First, the initial TJ motif is edited out to abolish activity in OSC cells. Second, a variety of S2-specific motifs are added, including GATA, ETS, SERBP, and AP-1 motifs, with one of the ETS sites overlapping where the initial TJ used to be. Together, these edits switch the specificity of the regulatory element from one cell line to the other.

## 5 Affinity catalogs reveal how Ledidi uses *cis*-regulatory logic to achieve precise outcomes

Although design methods are primarily used because the designs themselves are of practical utility, these methods can also be used scientifically to extract *cis-*regulatory rules learned by the models and to better understand relevant properties of the initial template sequences. We approach this by introducing the concept of an “affinity catalog”, where designs are produced for several steps across a range of target value strengths, and the edits made at each step can be compared. As a demonstration, we constructed fine-grained affinity catalogs for the SMYD3 promoter for three increasingly sophisticated modalities in human K562 cells: the binding of GATA2, chromatin accessibility, and transcription initiation, designed using BPNet, ChromBPNet, and ProCapNet models respectively.

For each modality, Ledidi successfully produced designs that are predicted to closely match the desired activity at each step (Fig 3A). We observed that this agreement diminished toward the edges of the catalogs, with the most notable disagreements occurring at very high desired levels of GATA2 binding. We initially hypothesized that this divergence arises because Ledidi cannot make designs more extreme than those observed in the genome; however, the 99th percentile of GATA2 model predictions is +0.992 greater than the original prediction and the maximum prediction +2.739 greater (the black bar), where divergence does not yet occur. Instead, this divergence is related to the balance between the input and output losses (Supplementary Fig 6). Because each edit imposes a cost, Ledidi begins with edits that individually yield the strongest effects. Once such edits have been exhausted, each step toward more extreme GATA2 binding requires, on average, more edits to achieve. Eventually, the number of edits required to increase predictions comprises a larger portion of the total loss than is made up for by more closely matching the desired output. This finding indicates that affinity catalogs can be useful in practice for setting the mixture weight correctly, to ensure that this divergence does not happen near where the target value is.

**Figure 3:**
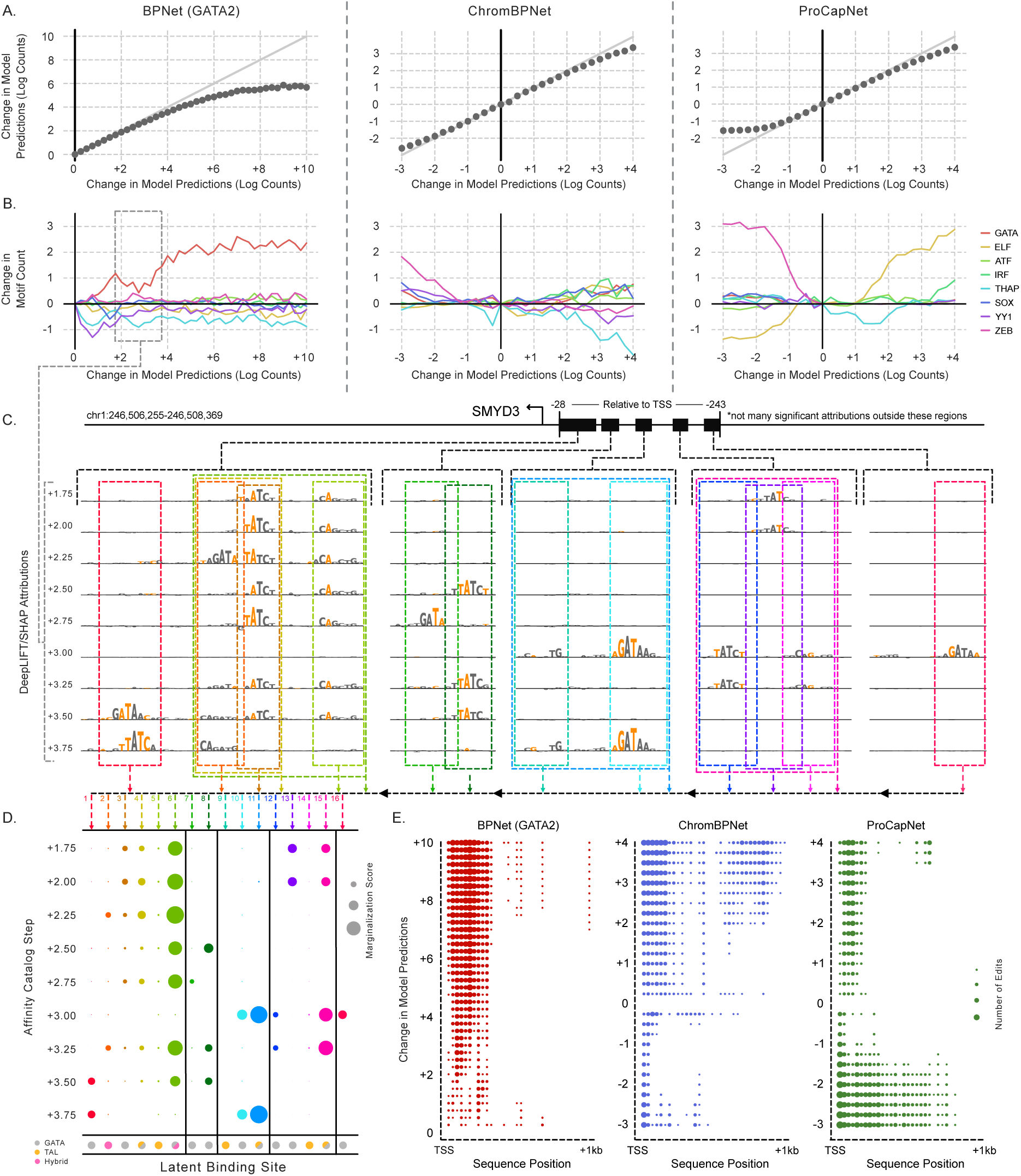
Affinity catalogs reveal subtle principles of *cis*-regulation. (A) Affinity catalogs for GATA2 binding, chromatin accessibility, and transcription initiation. Each model makes predictions for log counts, and the designs involve altering those predictions. At each step, design was performed 25 times and the predictions were averaged. (B) Change in motif composition across the catalogs, averaged across the 25 designs, with a shared x-axis as A. (C) Proposed edits and attributions for each step in a slice of the affinity catalog at several regions in the promoter region for SMYD3. Orange characters indicate those that have been edited, and colored boxes indicate the binding sites that are to be run through *in silico* marginalization. (D) A dot plot indicating how strongly each isolated binding site drives model predictions, quantitatively displaying how alternative patterns of editing latent binding sites achieves the specific outcome. (E) The average number of edits made at each 25 bp bin between the TSS and 1 kbp upstream of the promoter at each step of each of the three catalogs.

To better understand the edits made in these catalogs, we calculated the change in motif composition at each step (Fig 3B). An increasing number of GATA motifs was the primary trend in the GATA2 affinity catalog. The accessibility catalog involved multiple trends, with ZEB binding sites added to decrease accessibility and a mix of various binding sites being used to increase accessibility. In the transcription initiation catalog, ELF binding sites are clearly the primary driver of predictions, with Ledidi eliminating them to decrease predicted transcription initiation and adding them to increase it. Further, ZEB sites are also used to reduce transcription initiation, likely through their influence on accessibility, and THAP sites are depleted for only a portion of the catalog. By considering affinity catalogs in this manner, we can extract the motifs learned by ML models in a manner orthogonal to standard model interpretability methods.

Further inspection revealed that Ledidi was leveraging subtle properties of the *cis-*regulatory codes learned by these models to precisely match the desired output. For example, we observed in the GATA2 catalog a dip in the number of GATA motifs used in the designs between +1.75 and +3.75, with those at +2.75 using almost half the number of motifs, on average, as those at +1.75 (Fig 3B), despite model predictions still steadily increasing. Model attributions and the choice of edits at each step in this slice of the affinity catalog (Fig 3C) revealed that this dip in GATA motif count was possible because Ledidi was making edits to control motif spacing, presence of co-factors (namely TAL), and motif affinity. At the first visualized step, Ledidi introduced edits in an imperfect GATA motif (annotated as [a.a.] 3) near a TAL binding site (a.a. 5), and this imperfect GATA site is made stronger in the next step. In the third step, Ledidi makes edits in a second GATA motif (a.a. 2) in the opposite orientation at a sub-optimal spacing relative to the first one, and the upstream GATA site (a.a. 13) is not edited in. Interestingly, this second GATA motif (a.a. 2) is natively a weak TAL binding site and so, when left unedited and paired with an appropriately-spaced GATA site (a.a. 1), enhances the binding. As a demonstration of Ledidi’s robustness, many edits are consistently made across this slice of the catalog despite each run of Ledidi being independent. By considering each step, we can see how latent binding sites are alternatively used without necessarily needing more binding sites to achieve stronger outcomes.

By running *in silico* marginalizations (see Methods for details) for each of these latent binding sites and their surrounding contexts, we can quantify the influence of these editing choices (Fig 3D). As a clear example, the third block involves a TAL motif (a.a. 9) and a GATA motif (a.a. 10). Despite being weakly highlighted by attributions, this TAL motif itself does not contribute to predicted GATA binding, whereas the GATA site does, and the combination yields even stronger predicted binding (a.a. 11). By far the strongest sites were those that involved both a GATA and an appropriately spaced TAL motif, with individual GATA sites having a weaker score and TAL sites having almost no influence on predictions by themselves. However, subtle patterns were also at play, such as how the precise edits being made at one of the primary GATA sites (a.a. 3) led to different changes in predictions by itself and also when paired with the nearby TAL site, whose edit remains largely consistent across the steps. More broadly, these marginalizations can show us how alternative usage of these latent binding sites — and the specific edits at each site — can lead to precise target values being achieved.

We then characterized where Ledidi made edits for each of the three modalities (Fig 3E). We found that the GATA edits were primarily in the ∼300 bp window before the transcription start site, with this region being more densely edited as the target strength increases. Although many edits targeting chromatin accessibility and transcription initiation fell within this window as well, many edits were also made outside this window for those modalities. Edits were made upstream when increasing chromatin accessibility but not when decreasing it, whereas the opposite was true for transcription initiation. This suggests that the motifs driving chromatin accessibility are more concententrated toward the TSS, because making edits there disrupts accessibility, whereas the motifs driving transcription initiation are more dispersed and require editing a broader window to disrupt.

## 6 Designing affinity catalogs enables control over transcriptional dosage

Having shown that Ledidi can quickly design enhancers that exhibit cell type-specific expression *in cellulo*, we next investigated the extent to which Ledidi can quantitatively control the strength of the desired activity, and thus the transcriptional dosage. This is in contrast to our previously described enhancers, which were designed only to maximize cell type-specificity. Here, we designed three affinity catalogs using the same DeepSTARR models as before but replacing the min-gap loss with target values derived from the 33rd, 66th, 99th, and an interpolation of 50% stronger than the strongest enhancer according to genome-wide STARR-seq data in the target cell lines (see Methods for details). Two of these affinity catalogs were cell type-specific with increasing target values only in the on-target cell line, and the third catalog involved designing enhancers active in both cell lines. So as to better compare across settings, we used the same initial template sequences as in our previous experiments (Fig 4A).

**Figure 4:**
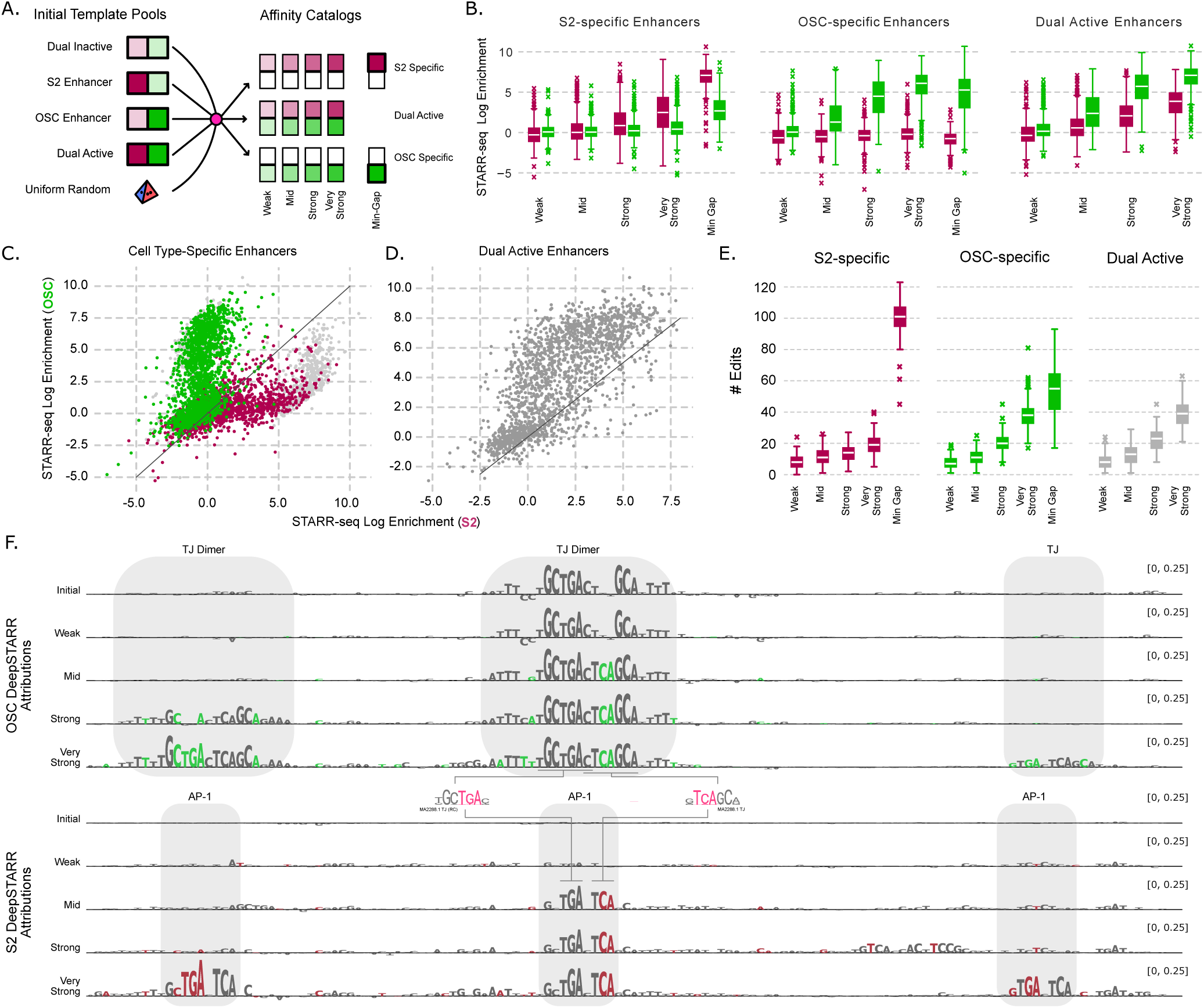
Designed affinity catalogs enable fine-grained control of gene dosage. (A) A cartoon demonstrating how genomic sequences from each pool are edited to induce various outcomes. Burgundy depicts results in S2 cells and green in OSCs. (B) STARR-seq enrichment in each cell type for each step in the affinity catalog, and for each target cell line(s). (C) STARR-seq enrichment for each designed regulatory element, with those targeting S2 in burgundy and those targeting OSC in green. The min gap-designed elements for either cell line is shown in light gray. (D) The same as C but for when the target is both cell lines. (E) A distribution of the number of edits made by Ledidi for each step in each catalog, given a total sequence length of 249. (F) Attributions from DeepSTARR models at each step for a region initially an OSC enhancer and designed to be active in both cell lines.

We found that the regulatory elements designed for these catalogs displayed a controllable range of cell type-specific transcriptional dosage (Fig 4B). Each step in each catalog exhibited stronger expression than the previous step in the on-target cell line while limiting the expression of the off-target cell line. The S2-specific catalog exhibited median enrichments of 0.82x, 1.00x, 1.81x, and 5.6x over controls in S2 cells, while the OSC-specific one exhibited 1.04x, 2.44x, 22.44x and 68.37x median enrichment over controls across the four steps in OSC cells. Neither catalog exhibited more than a 1.33x median enrichment in the off-target cell line at any step, and all of the steps in the OSC catalog were below 1x. Of particular note was that the strongest designed S2-specific enhancers in our catalog exhibited significantly weaker off-target activity (1.33x median enrichment) than the min-gap designed ones (6.35x median enrichment), exemplifying the balance between having the strongest on-target activity and minimizing off-target activity. When designed to be active in both cell lines, the elements exhibited stronger activity than the designed cell type-specific elements: the median S2 enrichment in the very strong dual enhancers was 2.61x higher than the median S2 enrichment in the very strong S2-specific elements, and the respective enrichment was 1.99x for OSC-specific elements.

When considered without grouping by activity bin, the designed elements in the cell type-specific catalogs exhibited a wide range of enrichments in the on-target cell line, and the dual-active elements largely exhibited activity in both cell lines, although they were somewhat stronger in OSCs than S2 cells (Fig 4C/D). In terms of activity, we observed that there was overlap between the elements in our S2 catalog and the min-gap designed elements, but those in our catalog avoided a population of elements that exhibited very strong S2-specific activity at the cost of higher activity in OSC as well, explaining our previous findings.

Interestingly, the elements in the affinity catalogs required far fewer edits than one might expect, with a median of 8/7 edits needed to design weak S2/OSC enhancers, respectively, and 19/38 to design the strongest enhancers. Far fewer edits were needed to design the strongest S2-specific enhancers in the catalog (19) than when using the min-gap loss (101). Fewer edits were needed in the OSC catalog even though the strongest enhancers exhibit stronger enrichments than the min-gap designed ones (Fig 4E). Together, these results demonstrate that Ledidi is capable of carefully designing edits that control both the specificity of the activity and the transcriptional dosage in the on-target cell line.

Next, we considered the specific edits made to a regulatory element that was initially an OSC-specific enhancer and was then edited to be active in both cell lines. Attributions from the DeepSTARR models revealed that a TJ motif drives the initial activity in OSC cells, and that this motif is edited to become a dimer in the stronger elements (Fig 4F). However, the precise positioning of the TJ instances, with the dimers overlapping by one nucleotide, creates an AP-1 motif within the TJ dimer itself. Because TJ motifs drive activity in OSC cells and AP-1 motifs drive activity in S2 cells, this nested motif drives activity in both cell lines. When tasked with designing very strong enhancers, we see that this AP-1/TJ motif is repeated on the flanks, demonstrating that it is a reproducible sequence feature that Ledidi is using, likely because it uses fewer edits than adding in two separate motif instances. When experimentally verified, we see that this element goes from exhibiting a 0.26x/4.70x enrichment in S2/OSC cells to 3.87x/7.59x enrichment in the very strong enhancers, confirming that this sequence feature acts in the anticipated manner.

## 7 Discussion

In this work, we introduced Ledidi, a DNA design method that uses predictive ML models to guide the design of edits to an initial template sequence. We first demonstrated Ledidi’s capabilities *in silico* by applying it out-of-the-box to dozens of models to design edits that increase TF binding, chromatin accessibility, transcription, and enhancer activity. We then tackled the more challenging problem of designing cell type-*specific* elements, and STARR-seq experiments confirmed that these elements exhibited strong and specific activity. Finally, we introduced the idea of an “affinity catalog”, where the design process is repeated across several steps of target strength, and we showed how these catalogs can be used for scientific exploration and to design elements that control transcriptional dosage in a cell type-specific manner.

A key conceptual difference between Ledidi and *de novo* design approaches is that Ledidi begins with an initial sequence and minimally edits it to induce desired characteristics. Focusing on edits has two practical consequences. First, Ledidi can be used for scientific investigation because the edits can be inspected to better understand the existing and latent *cis-*regulatory features of the initial template, as well as what the model is focusing on. Such investigation offers potential for an orthogonal class of model attribution algorithms that, rather than only explaining what drives predictions, can be used to illuminate what is *almost* there and provide a path for how to get there, e.g., how to activate an inactive sequence. Second, we anticipate that a component of the most successful designs will be choosing a good initial sequence, especially when the desired properties are complex and not fully captured by a single ML model. When the optimal choice is not immediately clear, we recommend using collections of public genomics datasets to find regions that already exhibit many of the desired characteristics [51]. However, when there truly is no reasonable template sequence, Ledidi still allows one to adopt a *de novo* approach by editing a randomly generated or randomly selected sequence.

A frequent challenge when designing regulatory elements is that one wants to precisely control multiple characteristics simultaneously, e.g., activity in multiple cell types. Ledidi naturally accommodates multi-term design objectives, where each term can come from a different output from the same model or even from separate models that have been trained in different settings. This point is important for several reasons. First, because models are usually developed with a specific goal in mind, the state-of-the-art model for one modality may not also make state-of-the-art predictions for all modalities one cares about. Second, when a new assay is developed or an old assay is applied in a new setting, it is likely infeasible to retrain and comprehensively evaluate massive models, such as Enformer, with this relatively small amount of additional data included. Instead, Ledidi offers the potential to train a light-weight specialist model for these new experiments to be used alongside the original massive model during design. Finally, even when a multi-output model makes predictions for all the modalities one requires for their designs, some outputs may be less accurate than others, even to the point where they cannot be trusted for design. In these settings, replacing the defective outputs with a light-weight specialized model may be necessary.

Although increasing model complexity generally also improves predictive performance, this complexity can also significantly slow down methods like Ledidi. For example, we found that BPNet using an A100 SXM4 40G GPU usually takes less than five seconds to design edits increasing TF binding for one sequence, but can take several minutes to do the same with Enformer, even when using predictions made for the same underlying experiment. This slowdown can come from two sources: using a model that considers sequences much longer than needed for your design task, and using a model with slow internal units such as transformers. Accordingly, we suggest that designers consider inference costs when choosing which model is best for their setting, and that model developers consider inference costs as a benchmark when proposing new models.

An important note is that model-based design methods such as Ledidi are only as good as the underlying oracles. We strongly recommend that the first step of design, whether using Ledidi or any other method, is to rigorously validate the ML models used. This validation should begin with checking predictive performance if that has not already been done, but should then continue to more sophisticated analyses such as considering *in silico* marginalizations and attributions to ensure that the model responds appropriately to biologically relevant sequence features. When the ML model relies on statistical artifacts, or the training data quality is so poor that overall performance measures are uninformative, then successful design is unlikely. Finally, after validating that the model is high quality and using it to design sequences, we recommend thorough *in silico* validation of the designs before using them experimentally. Such validation can include checking that the designs exhibit the desired characteristics using external models not used in the design process, checking motif composition, and applying other available domain knowledge, as we have done here. Fortunately, such manual validation of Ledidi’s outputs is made easier by only having to consider the influence of a small number of edits.

## 8 Methods

### 8.1 Ledidi

Ledidi is a DNA design method that rephrases the task of designing discrete edits to an initial template sequence as a continuous optimization problem using the straight-through estimator. This optimization problem has a loss that is a mixture of two terms: the first is the distance between the model output using the edited sequence *X* and the desired output *ȳ* (the output loss) and the second is the distance between the initial sequence *X*_0_ and the edited sequence (the input loss). More formally, the original sequence is a one-hot encoded matrix *X*_0_ ∈ 0, 1*^n,d^*, where *n* is the length of the sequence and *d* is the size of the alphabet; the edited sequence is, similarly, *X* ∈ 0, 1*^n,d^*; the model output using the edited sequence is *f* (*X*) ∈ ℝ*^k,l^*, where *k* is the length of the output sequence and *l* is the number of output measurements per position, and the desired output is *ȳ* ∈ ℝ*^k,l^*. The objective function for this optimization is

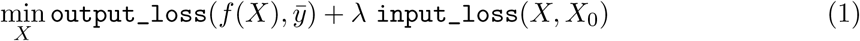

and we refer to the final edited sequence from Ledidi as *X̄*.

The input loss is weighted by the mixture weight *λ* to, when small, encourage a closer match to the desired output and, when large, encourage a closer match to the initial sequence. When *λ* is set to 0, Ledidi will design sequences that most closely match the desired output without considering the number of edits proposed, as in FastSeqProp [52]. Although any notion of distance could be used in each of the loss terms, we choose to use the L1 distance for the input loss corresponding to the number of edits made, and the L2 distance for the output loss:

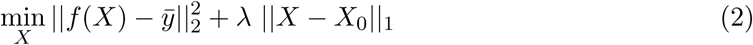

Unfortunately, Equation 2 is difficult to optimize because *X* is discrete, and continuous-valued gradients cannot be directly applied to one-hot encoded sequences. We overcome this challenge by carrying out the optimization with respect to a continuous weight matrix *W* ∈ ℝ*^n,d^*instead of over a one-hot encoded sequence. The matrix *W* is initialized to all zeros. At each step, we calculate *W* + *log*(*X*_0_ + *ɛ*) and use these values as logits such that each position in *n* is an independent Gumbel-softmax distribution over the alphabet *d* and a batch of one-hot encoded sequences are sampled from these distributions [53]. This sampling process is parameterized by a temperature *τ*, sometimes referred to as the “temperature”, that controls how close to the argmax the sampling is. As *τ* approaches 0, all sampled sequences resemble the most likely sequence in *W* + *log*(*X*_0_ + *ɛ*), and as *τ* approaches infinity, sampled sequences resemble those that would be drawn uniformly at random. We have found that Ledidi is robust to the precise value of *τ*, and so use a default value of 1. Because we are sampling discrete characters at each step, this process is referred to as the straight-through estimator [43].

With this new variable *W* we can solve an alternate form of Equation 1:

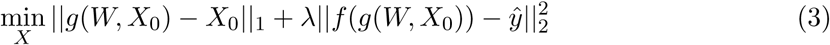

where *g* is the sampling process and *g*(*W, X*_0_) yields one-hot encoded sequences. This equation turns the process of designing edits into an optimization problem that can be solved with standard gradient descent methods. In this work, we use the Adam optimizer [54] with default settings except that the learning rate is set to 1.0.

Having an initial sequence enables several modifications to be made to the standard edit design task. First, one can specify a *mask* where edits cannot be made by setting the values of *W* to − inf at positions where edits are not allowed. This mask can be adapted to block specific categories of edits, e.g., not allowing edits to result in “G”s, or made more precise to not allow a specific single ““C” to become a “G” but still allowing it to become a “T”. Second, *in-painting* can be performed by removing characters or spans from the initial sequence and having Ledidi learn what character should appear at those positions. Another way of viewing in-painting is as an extension of the original design task, but with edits at certain positions able to be made for free, making their usage preferential. Finally, *priors* can be encoded in *W* with high values indicating the higher probability of that character being generated. These modifications can be seamlessly combined with each other and also with each other idea in this work without modification or additional burden on the user.

### 8.2 Training BPNet Models

Our evaluations involve many publicly available, pre-trained models, but we supplemented these with several BPNet models trained specifically for this work [26]. For each model, BAM and narrowPeak files were downloaded from the ENCODE Consortium data portal (https://www.encodeproject.org/). 5’ mapped read ends were pooled across replicates and then split across strands into two read count bigWig files using bam2bw (https://github.com/jmschrei/bam2bw). For the one chromatin accessibility data set, reads were not split across strands but instead kept as a single bigWig. Finally, we used bpnet-lite (https://github.com/jmschrei/bpnet-lite) to train BPNet models using standard settings (64 filters for TF binding and 256 filters for chromatin accessibility). These trained BPNet models, along with their training configurations, can be downloaded from https://zenodo.org/records/ 14604495.

### 8.3 Design of TF Binding Sites

As an initial validation of Ledidi, we designed edits that increased predictions across several models in fourteen design settings. A challenge with setting a standard design task across models is that these models were trained on data with different read depths and processing approaches, and make predictions for different outputs using different loss functions. For example, Beluga [45] predicts peaks as a classification task and only makes one prediction per output, and Enformer makes predictions for many 128bp bins for each output. These differences mean that there was not always a clear equivalent single value to set as Ledidi’s desired output. Accordingly, for models that predicted entire tracks (Borzoi and Enformer) we summed the signal across the output window, and for BPNet/ChromBPNet/ProCapNet models we use only the count head.

To account for differences in read depth and processing, we phrased the design task using percentiles, which would be agnostic to the underlying data distribution and comparable across settings. To get these percentiles, we chunked chr12 in hg38 into non-overlapping spans of the length expected for each model, removed all chunks that had *any* unknown characters, and made predictions using the model for the relevant task. From this distribution, we chose the 100 regions closest to the 50th percentile to use as our initial sequences, and tasked Ledidi with designing edits to bring them to the 99.5th percentile. Because the *λ* parameter is sensitive to the magnitude of the predictions, we set it to be

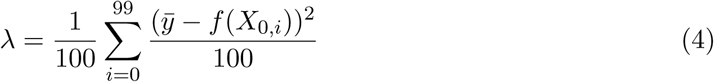

which is the average Euclidean distance between the initial and desired outputs divided by 100. This 100 value is unrelated to the choice of using 100 regions closest to the 50th percentile but, rather, corresponds to wanting each edit to reduce the loss by at least 1%.

This process was then repeated for the 46 design settings used subsequently to evaluate motif compositional changes.

### 8.4 Affinity Catalog Design

Affinity catalogs are collections of edits derived from running a design method repeatedly on a fine-grained range of target outcomes. These outcomes can involve increasing or decreasing model predictions, and include outcome values that fall well outside the training distribution for the ML model. Although one may naturally associate “affinity” with only TF binding, these catalogs can be designed for any characteristic that a ML model can make a prediction for. They are named such based on the intuition that the binding of TFs drives many important processes including accessibility and transcription, and, accordingly, affinity of TF binding still drives fine-grained changes in these higher-order characteristics. No special technique is needed to create an affinity catalog other than running the design step repeatedly. We designed affinity catalogs in three settings.

In Section 5 we designed *affinity catalogs* for GATA2 binding, chromatin accessibility, and transcription initiation at the SMYD3 promoter. These catalogs were designed using BPNet, ChromBPNet, and ProCapNet models in K562 respectively. The initial sequence was the SMYD3 promoter centered on the transcription start site. To simulate a realistic setting where one would not want to modify the gene body, and because the gene is on the negative strand, we masked out the first 1,057bp of sequence and only allowed edits in the second half of the sequence. For the GATA2 catalog our designs spanned an increase in predictions of 0 to 10.0 and a step size of 0.25. For the accessibility and transcription initiation catalog, our designs spanned a decrease in 3.0 to an increase of 4.0 and a step size of 0.25. We note that, for the three models used, these predicted values correspond to log fold change read/fragment counts in the region. We set *λ* using the same approach as when designing TF binding sites.

### 8.5 *In silico* marginalizations

In several evaluations we use *in silico* marginalization to quantify the influence that a motif has on model predictions. These evaluations are performed by implanting a provided motif into the center of a set of background regions and recording the difference in model predictions before and after the implantation [55]. In our evaluations, we used 1,000 background sequences that were constructed by dinucleotide shuffling the initial sequence being used for design. They are referred to as marginalizations because averaging across many background sequences marginalizes out the influence of the surrounding context and returns just the influence of implanting the motif.

### 8.6 STARR-seq Experiments

### 8.7 Building ladders

To make our measurements comparable to previous genome-wide STARR-seq experiments performed in S2 and OSC cells, we designed ladders composed of five bins of regulatory elements that exhibited various strengths in those experiments. These ladders contained the 50 strongest peaks, the 50 weakest peaks, and 50 elements selected at manually defined activity levels corresponding to roughly 25%, 50%, and 75% of the maximum strength (Supplementary Fig 7). This procedure was performed independently for each cell line without requiring that the elements in the ladders themselves were cell type-specific. DeepSTARR models were used to confirm that the bins were predicted to, on average exhbiit increasing strength.

#### 8.7.1 Generating libraries

300-bp oligonucleotides containing 249-bp candidate sequences were custom synthesized by TWIST and amplified as inserts for oligonucleotide STARR-seq plasmid libraries in two consecutive PCR steps. The first PCR was performed in the linear range with the primer pair (forward: ACACTCTTTCCCTACACGACGCTCTTCCGATCT, reverse: GTGACTG-GAGTTCAGACGTGTGCTCTTCCGATCT). In the second PCR amplification step, overhangs were added for Gibson cloning using the following primer pair (forward: TAGAGCATG-CACCGGACACTCTTTCCCTACACGACGCTCTTCCGATCT, reverse: GGCCGAATTCGTC-GAGTGACTGGAGTTCAGACGTGTGCTCTTCCGATCT. Amplified fragments were cloned using Gibson assembly (New England BioLabs, catalog no. E2611S) into the previously published STARR-seq plasmid containing the developmental Drosophila synthetic core promoter (DSCP (Pfeiffer et al 2008)) (Addgene 71499, Arnold et al. 2013). Oligonucleotide libraries were electroporated into MegaX DH10B electrocompetent bacteria (Thermofisher C640003) and grown in 2L LB-Amp (Luria-Bertani medium plus ampicillin, 100 µg/ml) and purified with a Qiagen Plasmid Plus Mega Kit (catalog no. 12981). The oligonucleotide library was then tagged with a unique molecular identifier (UMI), amplified and sequenced to derive an input count value for the representation of sequences in the plasmid library, as previously described (Neumayr et al. 2019).

#### 8.7.2 Cell culture

Drosophila S2 cells (Thermofisher R69007) grown in Schneider’s Insect medium (Gibco 21720001) with 10% fetal bovine serum (FBS) (Sigma-Aldrich, heat inactivated) and 1% P/S (Gibco 15140122) at 27°C and 0.4% CO2. OSC cells (obtained from DGRC, stock #288) were cultured in Shields and Sang M3 Insect Medium (Sigma #S8398) supplemented with 0.6 mg mL-1 glutathione (Sigma G6013), 10% FBS, 10 mU mL-1 insulin (Sigma I6634) and 10% fly extract. For electroporations the regular growth of cells was monitored, and cells were seeded at 3*10^^^6/ml density the day before electroporation. For each biological replicate, 50*10^^^6 S2 cells or 57*10^^^6 OSCs were resuspended in 100 µL of a 1:1 dilution of HyClone MaxCyte electroporation buffer and serum-free Schneider’s or M3 medium depending on the cell type and electroporated with 5 µg of the input libraries (see previous section) using the MaxCyte-STX system (‘Optimization 1’ protocol). Cells were then incubated in DNase I (Wothington Biochemical, 2000 U/mL) for 30 min and harvested for RNA extraction 24 h after addition of culture media for recovery and oligonucleotide library expression. RNA was extracted and processed following the previously described UMI-STARR-seq protocol (Neumayr et al. 2019).

#### 8.7.3 Illumina sequencing

Next-generation sequencing was performed at the VBCF NGS facility on a NovaSeq SP platform (150bp paired-end), following the manufacturer’s protocol, using standard Illumina i5 indexes as well as UMIs at the i7 index.

#### 8.7.4 STARR-seq data analysis

A custom index containing the corresponding sequences was generated using the buildindex function from the Rsubread R package (Liao et al 2019 version 2.10.0). STARR-seq paired-end reads were then aligned using the align function from the same package, with the following parameters: type = “dna”, unique = TRUE, maxMismatches = 2. Then, UMI sequences were retrieved and collapsed as previously described by de Almeida *et al* [36]. The replicates exhibited high correlation for each sample (Supplementary Fig 8), and the ladders exhibited the expected activities (Supplementary Fig 9).

#### 8.7.5 Data processing

This procedure resulted in three signal replicates for S2 and OSC and two input control replicates. For numeric stability, a pseudocount of 0.1 was added to the number of observations of each construct in each replicate. Each replicate was then individually read-depth normalized by dividing the number of observed reads for a construct by the total number of reads for that replicate, and these read-depth normalized replicates were averaged together. Constructs were removed if the total number of raw input reads summed across both replicates was fewer than 100, including constructs that were not measured in the input due to dropout. The enrichments presented in this paper are calculated as the averaged read-depth normalized experimental signal divided by the average read-depth normalized input signal, and are shown on a log2 scale.

### 8.8 Genome-wide UMI-STARR-seq data analysis, deep learning data preparation, and DeepSTARR modeling

#### 8.8.1 Genome-wide OSC UMI-STARR-seq data analysis

After initial quality checks using fastqc, the OSC STARR-seq reads were trimmed using trim galore (version 0.6.0) with default parameters and aligned to the dm3 version of the Drosophila genome using bowtie2 (version 2.3.5.1) with default parameters. Aligned reads with a mapping quality ¿= 20 were then UMI-collapsed (tolerating for one mismatch), and peaks were called using MACS2 (version 2.2.5) with the following parameters: -g dm –keep-dup all -f BED –nomodel –extsize 200 –SPMR -B. The two biological replicates were processed separately but pooled for further deep learning data preparation.

#### 8.8.2 Deep learning data preparation

The Drosophila genome was binned into 249-bp bins with a 50-bp stride. The chromosomes U, Uextra, as well as the mitochondrial genome, were excluded for further processing. All windows overlapping the OSC STARR-seq peaks and a set of inactive windows (non-peak regions) were selected. We augmented the dataset by including the reverse complement for every sequence. Thus, for training and evaluating the OSC model, we ended up with 247,290 sequences (494,580 sequences post-augmentation). For the S2 model, we downloaded the training, validation, and test sets from zenodo (https://zenodo.org/records/5502060).

#### 8.8.3 Training DeepSTARR models

We trained the DeepSTARR (de Almeida et al., 2022) model on one-hot-encoded DNA sequences and predicted enhancer activity as the log2 enrichment over input. Both OSC and S2 models were evaluated on the first (validation) and second (testing) half of chromosome 2R. The models were trained in PyTorch (version 2.5.1).

### 8.9 Data Availability

The raw sequencing data are available from GEO (https://www.ncbi.nlm.nih.gov/geo/) under accession number GSE312234.

### 8.10 Code Availability

The Ledidi code is free and open source under an Apache v2.0 License at https://github. com/jmschrei/ledidi. Tutorials, narrative documentation, and an API reference are available at https://ledidi.readthedocs.io/en/latest/.

## Acknowledgements

We thank Vincent Loubiere, Nikolaus Mandlburger, and Jay X. J. Luo for discussions, help, and feedback, and Julius Brennecke for support. FKL is supported by an ESPRIT fellowship of the Austrian Science Fund (10.55776/ESP9305624) and MH by a Boehringer Ingelheim Fonds PhD fellowship. BR is supported by a DOC Fellowship from the Austrian Academy of Sciences. Research in the Stark group is supported by the Austrian Science Fund (10.55776/P36971, 10.55776/PAT3564423) and the WWTF (10.47379/LS24012). Research at the Institute of Molecular Pathology (IMP) is supported by Boehringer Ingelheim GmbH and the Austrian Research Promotion Agency (FFG, FO999902549). For the purpose of Open Access, the authors have applied a CC BY public copyright license to any Author Accepted Manuscript (AAM) version arising from this submission. The computational results presented were obtained using the CLIP cluster (https://clip.science). Next-generation sequencing was performed by the Vienna Biocenter Core Facilities GmbH (VBCF) Next-Generation Sequencing Unit.

---

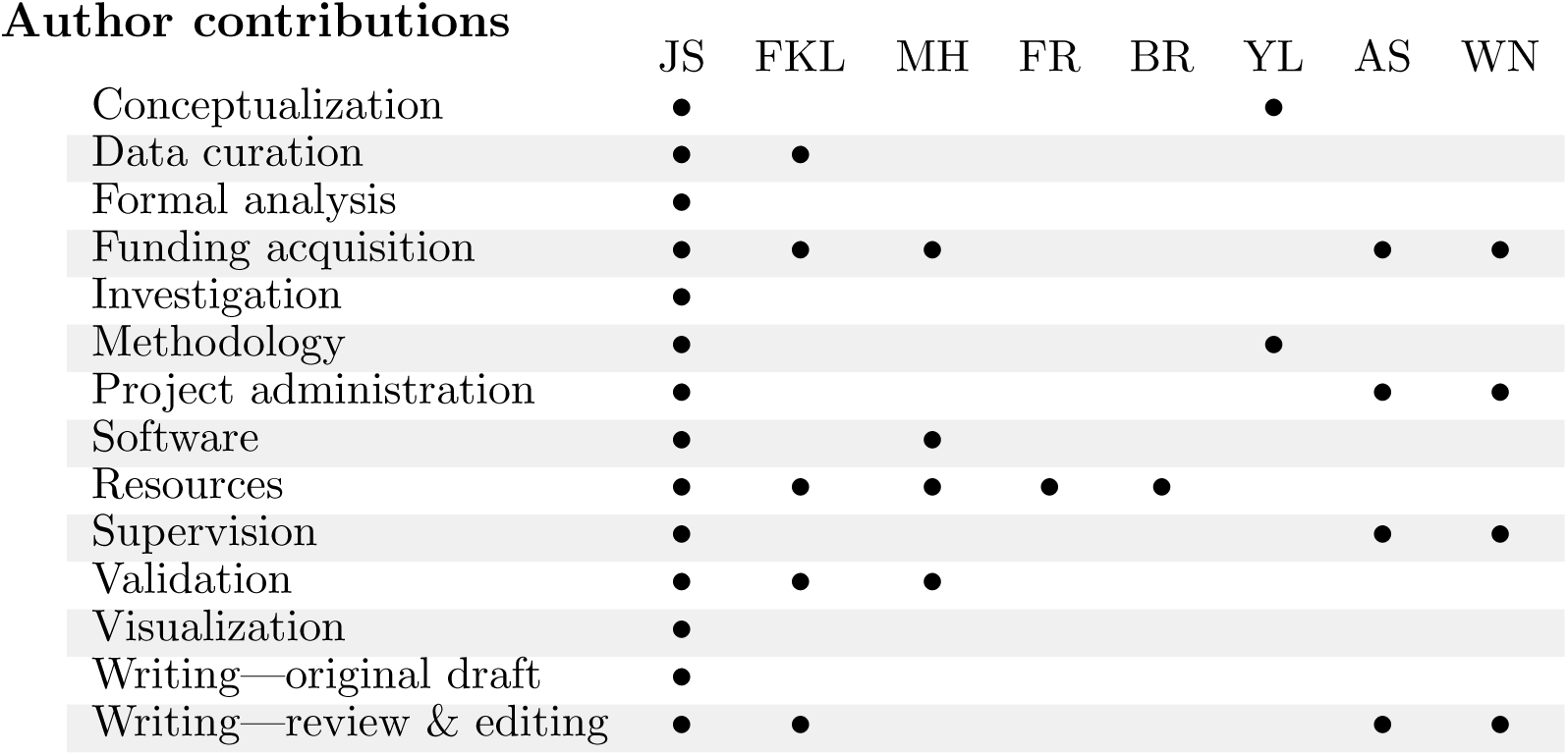

## Corresponding Author

Correspondance to Jacob Schreiber.

## Supplementary Materials

### Supplementary Tables

**Supplementary Table 1:** Metadata of models used for initial design. The type of machine learning model, assay the model was trained on, target of the assay (if applicable), output target used when the models are multi-task, and length of the input sequences used for each of the fourteen design settings. All design settings are in human K562 except for DeepSTARR which is in *Drosophila melanogaster* S2 cells.

| Model | Assay Type | Target | Target Index | Input Length |
| --- | --- | --- | --- | --- |
| BPNet | TF ChIP-seq | GATA2 |  | 2114 |
| BPNet | TF ChIP-seq | E2F6 |  | 2114 |
| Beluga | TF ChIP-seq | CTCF | 338 | 2000 |
| Beluga | TF ChIP-seq | MAX | 636 | 2000 |
| BPNet | ATAC-seq |  |  | 2114 |
| ChromBPNet | ATAC-seq |  |  | 2114 |
| Enformer | DNase-seq |  | 625 | 2000 |
| Sei | DNase-seq |  | 1490 | 4096 |
| ProCapNet | ProCap |  |  | 2114 |
| Puffin | ProCap |  | 5 | 1000 |
| Enformer | CAGE |  | 5111 | 2000 |
| Borzoi | CAGE |  | 306 | 2048 |
| Malinois | MPRA |  | 0 | 600 |
| DeepSTARR | STARR-seq |  | 0 | 249 |

### Supplementary Figures

**Supplementary Figure 1:**
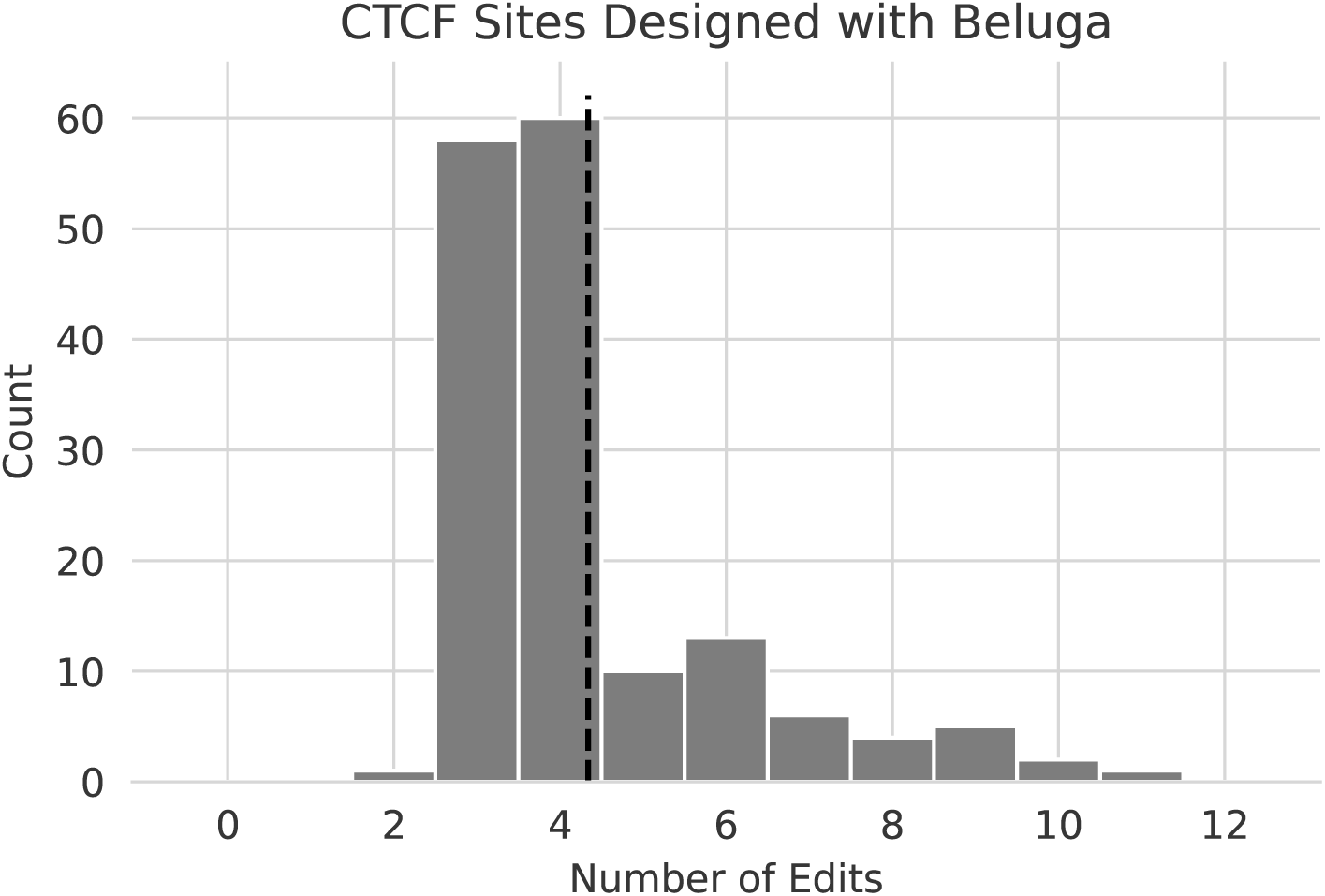
Number of edits needed to design CTCF binding sites. A distribution of the number of edits that Ledidi makes across 160 sequences to raise median CTCF predictions (indicative of no binding activity) to the 99.5th percentile (indicative of strong binding activity) using the Beluga/DeepSEA model. An average of 4.34 edits are made (the black dashed line). As the receptive field of this model is 2 kbp, this corresponds to changing only ∼ 0.2% of the sequence.

**Supplementary Figure 2:**
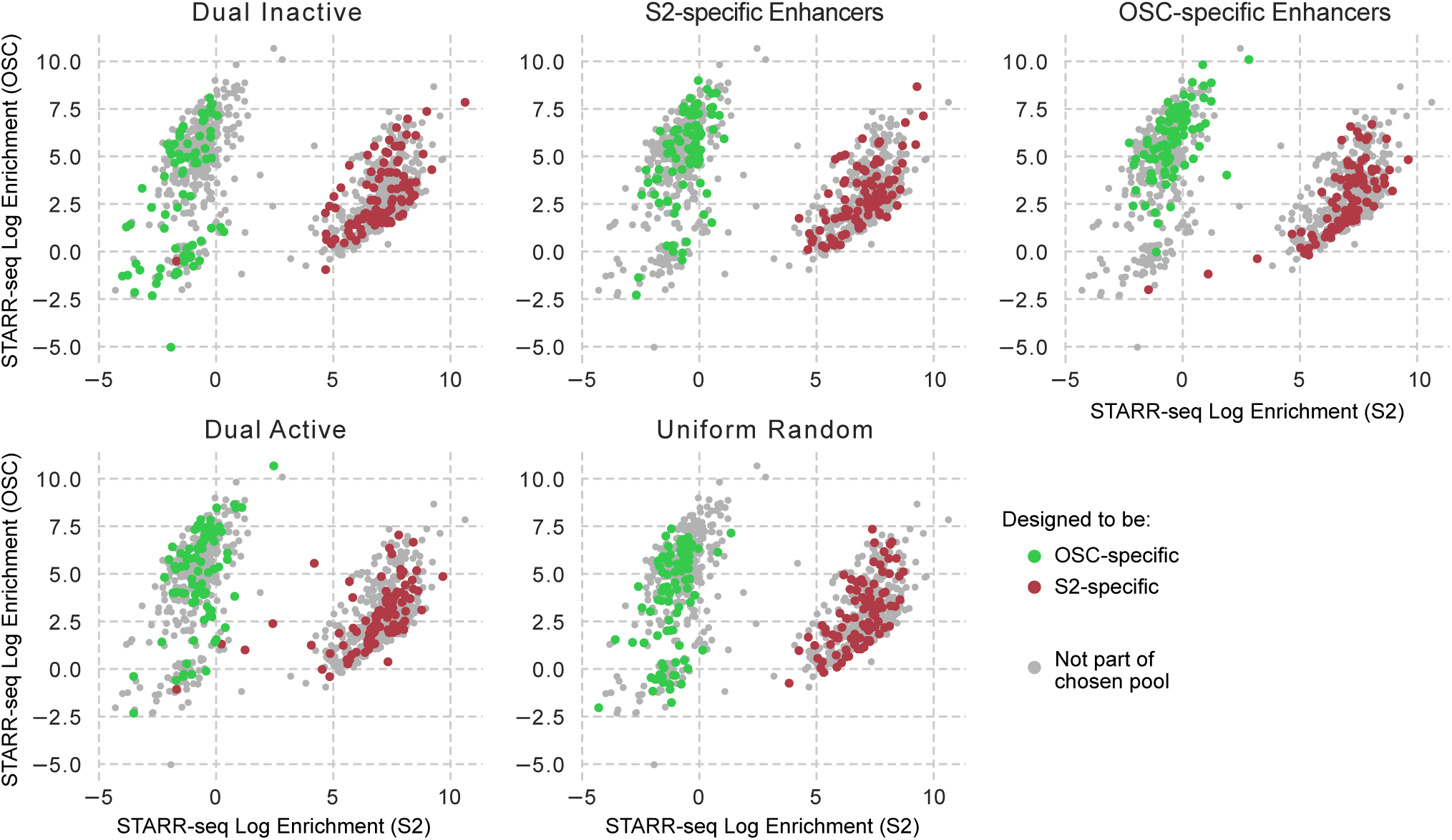
Cell type-specific enhancer designs stratified by initial pool. The STARR-seq enrichments for each designed element stratified by the initial pool, with the colored dots indicating that the enhancer is in the specified pools, the color corresponding to the target cell type, and the gray dots showing the complete set of designed enhancers.

**Supplementary Figure 3:**
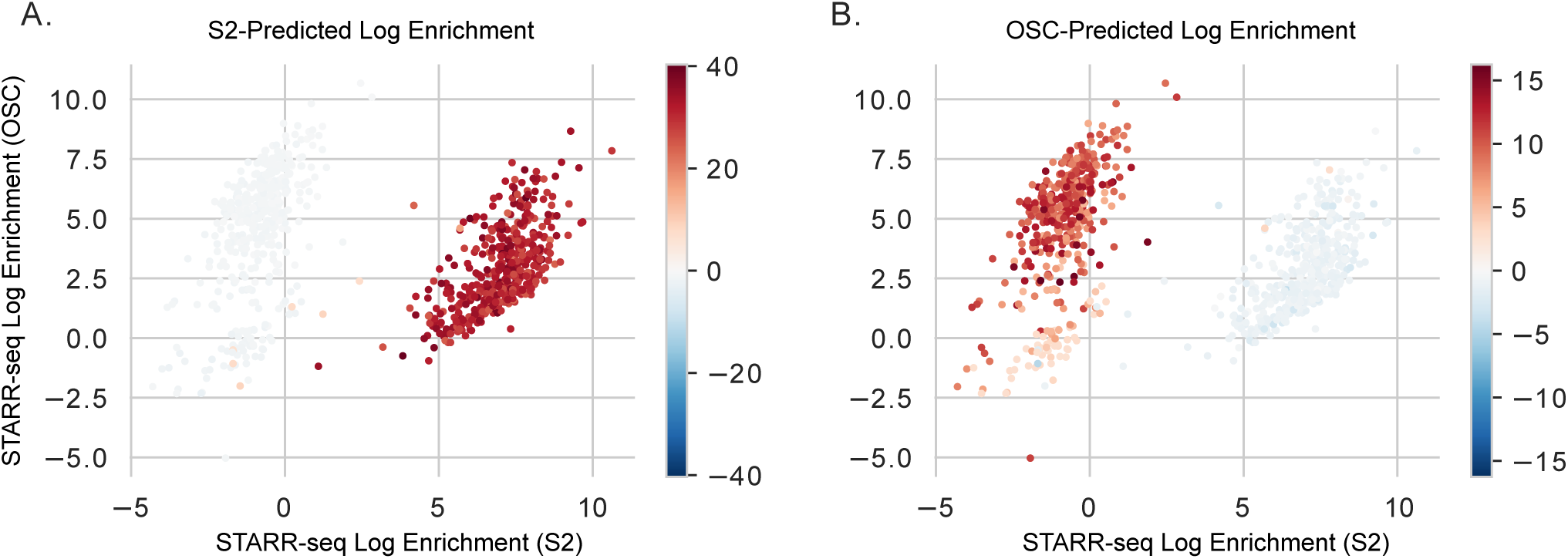
DeepSTARR predictions for the min gap-designed enhancers. (A) A scatterplot of the designed regulatory elements, with the positioning based on STARR-seq enrichments and the coloring based on the S2 DeepSTARR predictions. (B) The same, except using the OSC DeepSTARR model for predictions.

**Supplementary Figure 4:**
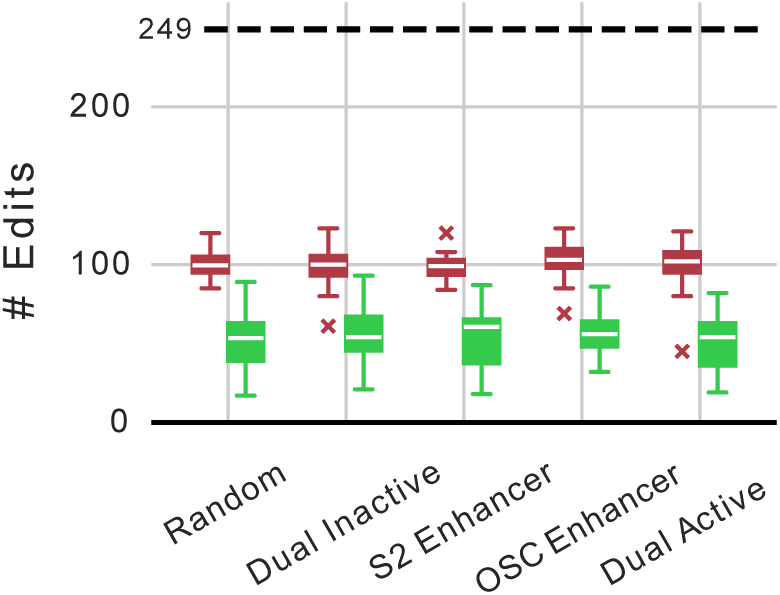
Number of edits made to design cell type-specific enhancers. A distribution of the number of edits made by Ledidi when using the min-gap loss to create cell type-specific enhancers, stratified by the pool of the initial template sequences.

**Supplementary Figure 5:**
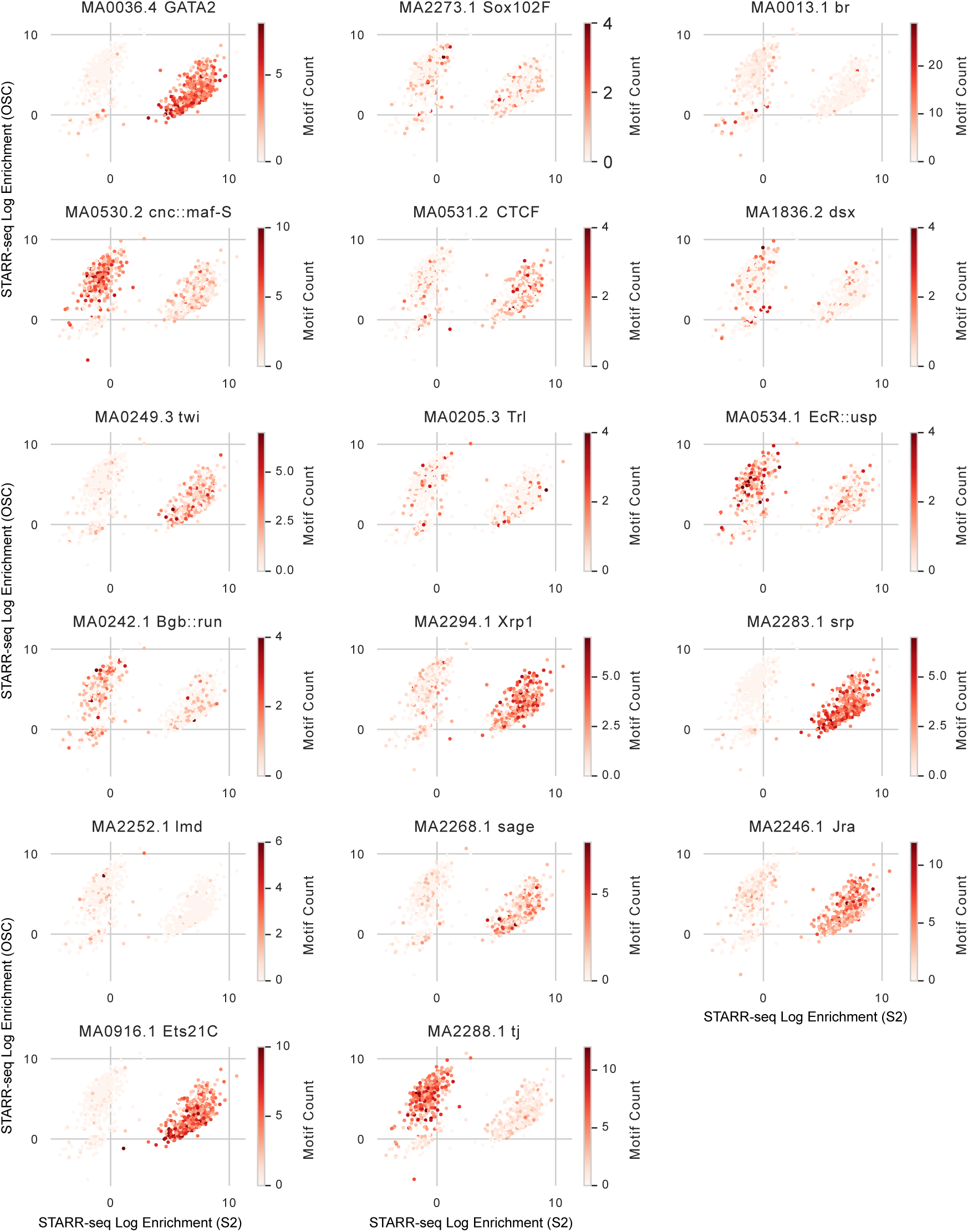
Motif counts in each design. A count of the number of a selection of motifs in each designed regulatory element when using the min-gap loss. Most motifs are present in either zero or one cell types, but not both, consistent with the task of designing cell type-specific activity.

**Supplementary Figure 6:**
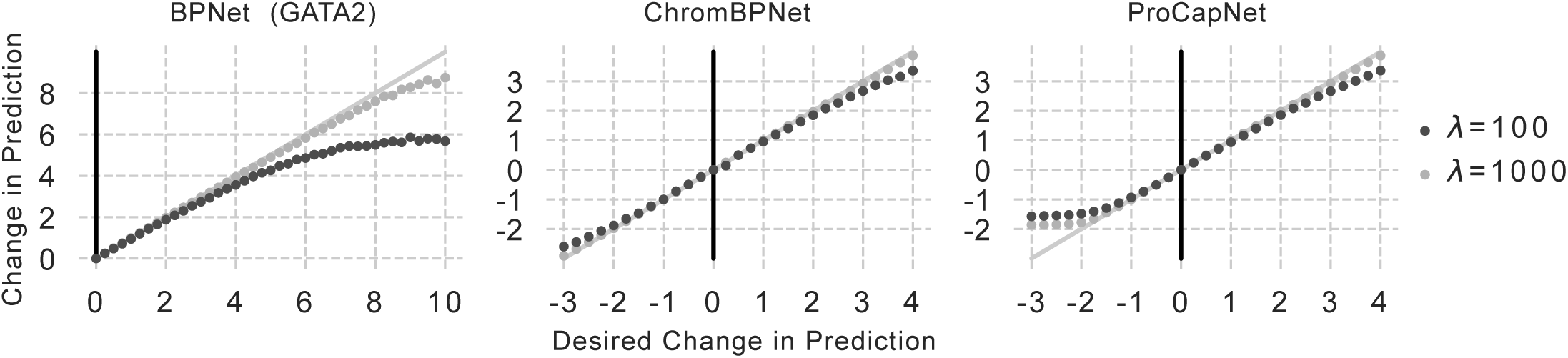
Divergence in affinity catalogs is explained by edit loss weight. We explored the cause of the divergence between the desired change in predictions and actual changes in predictions when designing affinity catalogs by reducing the penalty for edits by an order of magnitude (light gray). We observed significantly less divergence in all three tasks, though we note that the divergence does appear to still happen eventually. This is consistent with the trade-off between the input and output losses causing the divergence.

**Supplementary Figure 7:**
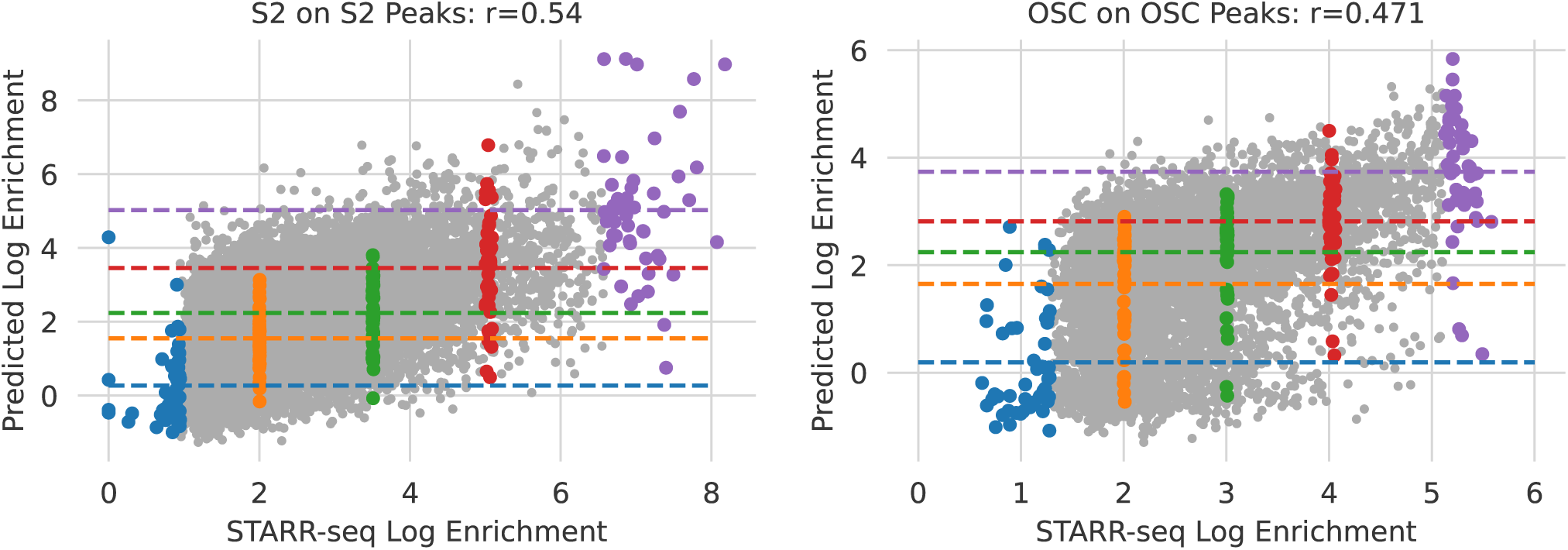
Choosing elements for STARR-seq ladders. As a control for our experiment validating the Ledidi-designed elements, we created ladders from STARR-seq peaks of various strengths according to previous STARR-seq experiments in the two relevant cell lines. Our ladders included the 50 strongest elements, the 50 weakest elements, and then 50 elements at manually defined levels evenly separating the range of activity. The chosen elements are colored according to the bin they were chosen for, with elements chosen for the S2 ladder on the left and those chosen for the OSC ladder on the right.

**Supplementary Figure 8:**
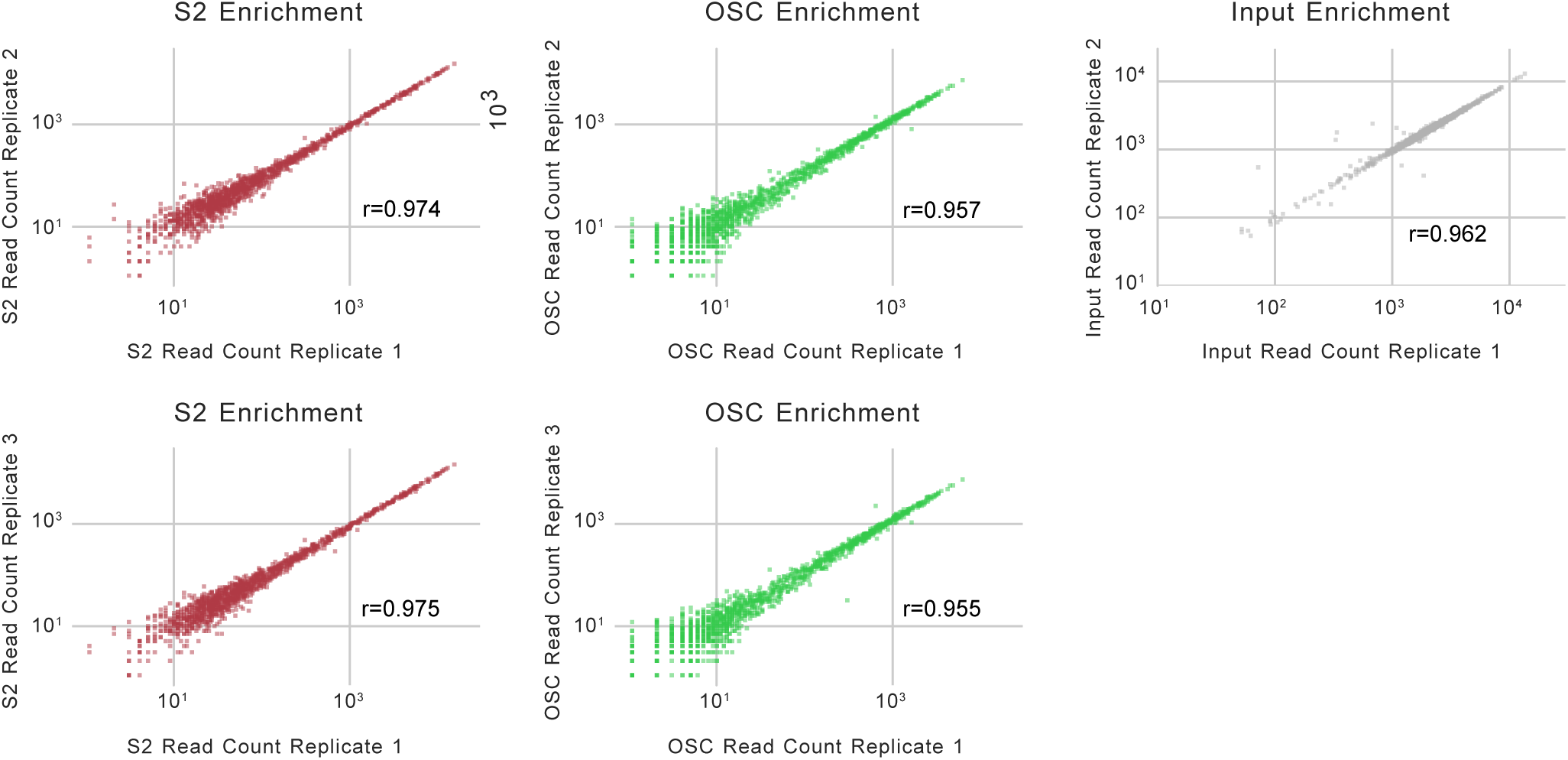
Replicate correlation for each sample. Three replicates were performed in S2 and OSC cell lines and two replicates were performed for the input control. Pearson correlations were calculated between replicates 1 and replicates 2/3 in S2/OSC cells, and the two replicates for the input control.

**Supplementary Figure 9:**
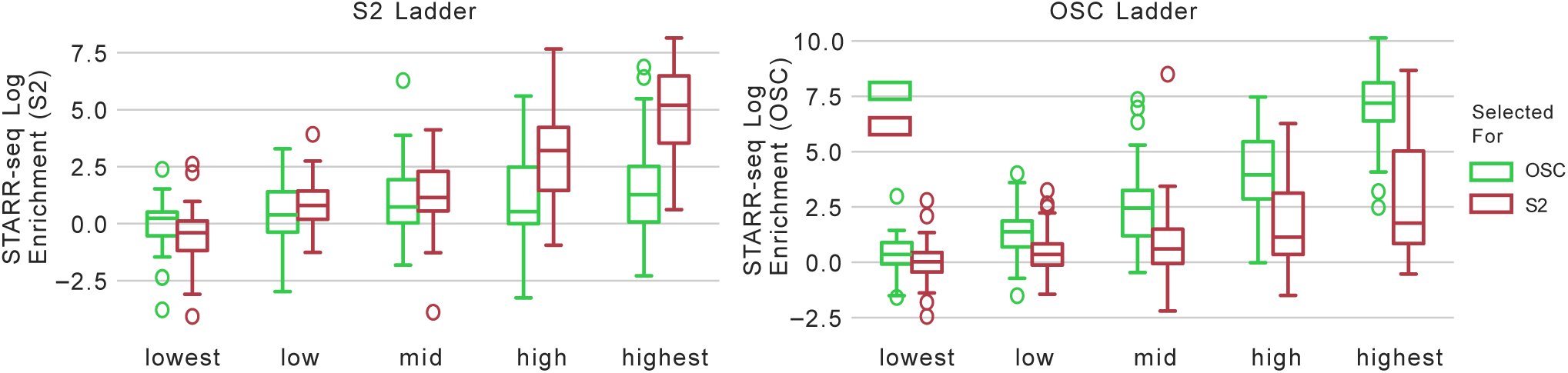
Activity for ladders in our STARR-seq experiment. The STARR-seq enrichments for our chosen ladder elements in both cell lines. These elements were not explicitly chosen to be *specific* to their cell line and, consequently, we see increasing activity in both cell lines in general but much stronger increases in the cell line the ladder was chosen for.

